# Direct and indirect effects of maternal, paternal, and offspring genotypes: Trio-GCTA

**DOI:** 10.1101/2020.05.15.097840

**Authors:** Espen Moen Eilertsen, Eshim Shahid Jami, Tom A. McAdams, Laurie J. Hannigan, Alexandra S. Havdahl, Per Minor Magnus, David M. Evans, Eivind Ystrom

## Abstract

Indirect genetic effects from relatives may result in misleading quantifications of heritability, but can also be of interest in their own right. In this paper we propose Trio-GCTA, a model for separating direct and indirect genetic effects when genome-wide single nucleotide polymorphism data have been collected from parent-offspring trios. The model is applicable to phenotypes obtained from any of the family members. We discuss appropriate parameter interpretations and apply the method to four exemplar phenotypes; offspring birth weight, offspring temperament, maternal relationship satisfaction, and paternal body-mass index, using real data from the Norwegian Mother, Father and Child Cohort Study (MoBa).

## Introduction

Most human traits exhibit some degree of heritability (Polderman et al., 2015). Some phenotypes are characteristics not only of individuals, but also depend on the influence of other individuals. While *direct* genetic effects refer to how the phenotype of an individual depends on their own genotype, *indirect* genetic effects refer to how it depends on the genotypes of others (McAdam, Garant, & Wilson, 2014). In this paper we describe a model for separating direct genetic effects from the indirect genetic effects of family members when genome-wide single nucleotide polymorphism (SNP) data have been collected from parent-offspring trios.

As parents transmit half their complement chromosomes to their children, the genomes of parents and offspring are correlated. Because the same genetic variants can have both direct and indirect effects, failing to account for the indirect genetic effects of relatives when attempting to measure heritability can result in misleading quantifications of the importance of direct genetic effects (Eaves, Pourcain, Smith, York, & Evans, 2014; Young, Benonisdottir, Przeworski, & Kong, 2019).

Indirect genetic effects can also be of interest in their own right. With respect to the focal individual (i.e., the individual whose phenotype is the focus of study), indirect genetic effects are part of the environment and may be of great interest when trying to understand causes of individual differences. In this paper we are concerned with indirect genetic effects underlying intra-familial dynamics. This can include instances where heritable characteristics of parents affect offspring development. For example, maternal influence offspring health through the intrauterine environment (Evans, Moen, Hwang, Lawlor, & Warrington, 2019), or where parents affect offspring development by providing an advantageous rearing environment. It also includes instances where heritable characteristics of the offspring evoke responses in their parents. For example, when child behavior influences the mental wellbeing of their parents.

The quantitative genetics literature distinguishes between two approaches to modelling indirect genetic effects. Trait-based models specify indirect genetic effects on the phenotype of the focal individual mediated by the phenotypes of other individuals. Variance-partitioning models avoid specification of the phenotypes that underlie the indirect genetic effects, instead quantifying the total contributions from these effects while being agnostic as to the underlying mechanisms (Bijma, 2014).

The emergence of large-scale genotype data in population-based cohorts has provided new opportunities for developing methods to separate direct and indirect genetic effects. This was leveraged by Eaves et al. (2014) who proposed a variance-partitioning method for separating indirect maternal genetic effects from direct effects with respect to an offspring phenotype, relying on genome-wide SNP data from mother-offspring pairs. In the current manuscript we extend the work of Eaves et al. (2014) to separate direct and indirect genetics effects within parent-offspring trios. We discuss alternative interpretations of variance components depending on the role of the focal individual, useful restricted model specifications and apply the method to four etiologically diverse exemplar phenotypes (offspring birthweight, offspring temperament, maternal partner relationship satisfaction and paternal body mass index) using real data from the Norwegian Mother, Father and Child Cohort Study (Magnus et al., 2016).

## Model formulation

Yang et al. (2010) introduced a method for quantifying additive genetic variance contributions from all measured SNPs using a linear mixed effects model. Extensions of this methodology include formulations for quantifying dominance genetic effects (Zhu et al., 2015), gene-environment interactions (Yang, Lee, Goddard, & Visscher, 2013), parent-of-origin effects (Laurin et al., 2018), maternal effects (Eaves et al., 2014) and avoiding bias from environmental effects (Young et al., 2018). The current approach (Trio-GCTA) uses parent-offspring trios to quantify the importance of direct and indirect genetic effects within the nuclear family. We refer to the individual whose phenotype is under study as the focal individual, noting that the method is applicable regardless of who is the “owner” of the phenotype.

In order to formulate a model for direct and indirect genetic effects, we assume that phenotypic measures have been obtained from a focal individual in *K* parent-offspring trios, and that genotypes for the same *M* SNPs are available for all individuals. We represent the three *K* × *M* matrices of maternal, paternal and offspring standardized genotype dosages (Zhu et al., 2015) by **Z**_*m*_, **Z**_*p*_ and **Z**_*o*_, respectively, arranged so that row *k* corresponds to the same parent-offspring trio. A linear model for the phenotypes can then be formulated as

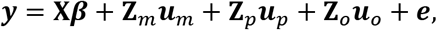

where ***y*** is a *K* × 1 vector of continuous phenotypes, **X** is a *K* × *P* matrix of measured covariates with *P* × 1 vector of coefficients ***β***, ***u***_*m*_, ***u***_*p*_ and ***u***_*o*_ are *M* × 1 vectors of additive random genetic effects associated with the maternal, paternal and offspring standardized genotype dosages, respectively, and *e* is a *K* × 1 vector of residual effects.

The genetic and residual effects are assumed to follow a multivariate normal distribution, where the different types of genetic effects may be dependent but individual SNP effects are independent. The residual effects are assumed to be independent of the genetic effects and across individuals

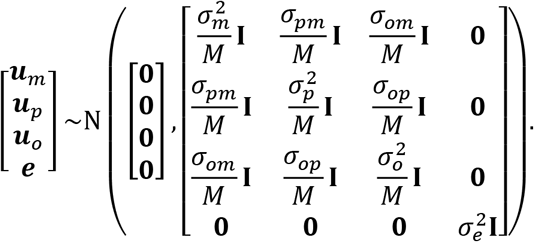

The expected covariance structure of the phenotype across all individuals is given by:

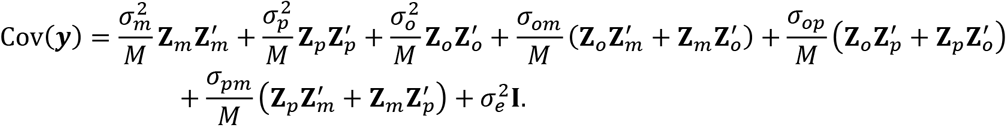

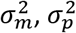 and 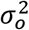 are the variances of the maternal, paternal and offspring genetic effects, respectively, *σ*_*om*_ is the covariance between the offspring and maternal genetic effects, *σ*_*op*_ is the covariance between the offspring and paternal genetic effects, *σ*_*pm*_ is the covariance between the paternal and maternal genetic effects and 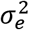 is the residual variance. When mating is random, the covariance between the maternal and paternal effects are not expected to contribute to the variance of the phenotype and the total variance decomposition is therefore

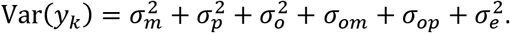

Depending on the role of the focal individual, the model parameters have different interpretations. If it is an aspect of the offspring phenotype that is under study, 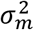 and 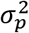 corresponds to variance attributable to indirect genetic maternal and paternal effects, respectively, whereas 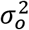 is the variance due to direct genetic effects. The components *σ*_*om*_ and *σ*_*op*_ are the covariances between the direct offspring genetic effect and the indirect maternal and paternal genetic effects, respectively. These parameters quantify the extent to which the same variants contribute to direct and indirect genetic effects. With respect to the offspring, the maternal and paternal genetic effects form part of the environment so these covariance terms may therefore also be interpreted as measuring variability due to gene-environment correlations. The component *σ*_*pm*_ is the covariance between the indirect maternal and paternal effects and is a measure of the extent to which the same variants contribute to indirect genetic effects. Sex-dependent expression of genetic effects has been studied with respect to a variety of phenotypes using family designs (Neale & Cardon, 2013). A weak correlation between maternal and paternal effects would indicate a qualitative sex difference, wherein mothers and fathers influence their offspring through different heritable traits (alternatively it could be that ostensibly the same trait is under the influence of different genetic factors when expressed in mothers and fathers). A correlation of unity but different magnitude between the maternal and paternal effect would indicate a quantitative sex difference, wherein mothers and fathers influence the offspring by the same heritable traits, but to a quantitatively different extent. Sex-dependent expression of parental effects can therefore potentially reveal insights into differences in maternal and paternal effects on the offspring. 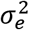 is the residual variance of the phenotype.

If it is an aspect of a maternal phenotype that is under study, 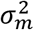 is the variance due to direct genetic effects, whereas 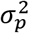 and 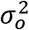 measure variability due to indirect genetic effects. The paternal and offspring genetic effects are environmental from the perspective of the mother. Although the underlying mechanisms may be distinct, a maternal phenotype may depend on interactions with both their partner and offspring. *σ*_*pm*_ and *σ*_*om*_ are the covariance between the direct maternal genetic effect, and the indirect paternal and offspring genetic effects, respectively. If the same genetic variants contribute to direct and indirect genetic effects, these covariance terms are expected to differ from zero. Assuming that mating is random, a genetic correlation between the direct maternal and indirect paternal effect is not expected to affect the phenotypic variance, because maternal and paternal genotypes are independent. However, as the offspring and maternal genotypes are correlated, a genetic correlation between the direct maternal and indirect offspring effect implies a gene-environment correlation that will either increase or decrease the phenotypic variance depending on the sign of *σ*_*om*_. *σ*_*op*_ is the covariance between the indirect paternal and offspring effects and is a measure of the extent to which the same additive genetic effects contribute to the indirect genetic effects. 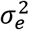 is the residual variance of the phenotype. These interpretations are conversely the same if it is a paternal phenotype that is under study.

## Special cases

Several other models of potential interest can be obtained as special cases of the general model described above. Young et al. (2018) introduced relatedness disequilibrium regression (RDR) as a method to avoid environmental bias in heritability estimates by modelling parental genetic nurturing effects in addition the direct genetic effects. The RDR model can be specified by setting 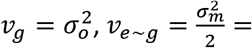 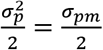 and 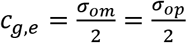, where *v*_*g*_ is the variance due to direct genetic effects, *v*_*e*~*g*_ is the variance due to parental genetic effects and *c*_*g*,*e*_ is the covariance between the direct and the parental genetic effects. Therefore, the RDR model can also be seen as assuming the maternal and paternal genetic effects are the same and of equal magnitude. If maternal or paternal effects are not of specific interest on their own, this will likely be a more effective way of accounting for indirect parental effects, as only four variance parameters are required compared to seven under the general model. Eaves et al. (2014) proposed a method (M-GCTA) for jointly estimating the variance explained by direct genetic effects, indirect maternal genetic effects and their covariance with respect to an offspring phenotype. The M-GCTA model can be obtained with the constraints 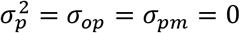. For many research questions, especially those related to pre- and peri-natal phenotypes, this may be a sufficient model. The original GCTA model (Yang, Lee, Goddard, & Visscher, 2011) can be obtained by omitting all indirect genetic effects from the model.

In the applications below we explore further interpretations of the model parameters when the focal individual has different roles. In the supplementary material we provide a simulation study demonstrating that parameters can be recovered when a trait is generated as a function of correlated direct and indirect genetic effects.

## Applications

We applied the Trio-GCTA method to a set of phenotypes measured in parent-offspring trios participating in the Norwegian Mother, Father and Child Cohort Study (MoBa) (Magnus et al., 2016). MoBa is a population-based pregnancy cohort study conducted by the Norwegian Institute of Public Health. Participants were recruited from all over Norway from 1999-2008. The women consented to participation in 41% of the pregnancies. The cohort comprises 114,500 children, 95,200 mothers and 75,200 fathers. The current study is based on version 11 of the quality-assured data files. Information was also obtained via a linkage to The Medical Birth Registry (MBR), a national health registry containing information about all births in Norway.

Blood samples were obtained from both parents during pregnancy and from mothers and children (umbilical cord) at birth. The project Better Health by Harvesting Biobanks (HARVEST) sampled 11,000 parent-offspring trios for genotyping from MoBa’s biobank at random. Genotyping was performed using llumina HumanCoreExome-12 v.1.1 and HumanCoreExome-24 v.1.0 arrays. The pre-imputation quality control and imputation procedure is described in Helgeland et al. (2019). Post-imputation, four and a half million SNPs with imputation info score > 0.9 and minor allele frequency > 0.05 were retained for analyses.

Example 1: Birth weight (offspring phenotype)

Both offspring and maternal genes are likely to be involved in determining birth weight as the intrauterine environment is provided by the mother. Both traditional family (Lunde, Melve, Gjessing, Skjærven, & Irgens, 2007; Magnus, 1984) and molecular genetic designs (Warrington et al., 2019) have previously indicated substantial portions of variance in birth weight determined by both direct offspring and indirect maternal genetic effects. We applied the current method to birth weight measures from 8154 trios in order to obtain a comparison to previous findings. The current method further allows the correlation between maternal and offspring genetic effects to be estimated.

Example 2: Infant temperament (offspring phenotype)

The Infant Characteristics Questionnaire was developed for assessing parental perceptions of infant difficult temperament (Bates, Freeland, & Lounsbury, 1979). Using a twin design, Silberg et al. (2005) found that they could account for as much as 75% of the variability in difficult temperament with direct additive genetic effects. Prenatal maternal characteristics such as depression and anxiety have also been suggested to affect infant temperament, but findings are mixed (Erickson, Gartstein, & Dotson, 2017). The importance of such prenatal factors can possibly be measured via maternal genetic effects. We analyzed a summated score of maternal reports to 10 items related to infant difficult temperament at age 6 months from 7291 trios.

Example 3: Relationship satisfaction (maternal phenotype)

Maternal reports of relationship satisfaction between mothers and fathers have been found to decrease on average following the birth of a child (Dyrdal, Røysamb, Nes, & Vittersø, 2011). A possible explanation for this decrease is that relationship satisfactions to some degree depend on aspects of the infant phenotype. We therefore investigated whether maternal reports of relationship satisfaction six months after birth are influenced by offspring genotype. Measures of relationship satisfaction were obtained by summation of the ten items comprising the Relationship Satisfaction scale (Røysamb, Vittersø, & Tambs, 2014) in 7187 trios.

Example 4: Body mass index (paternal phenotype)

Body mass index (BMI) in adulthood has both genetic and environmental components of causation. Yang et al. (2015) found that 27% of variability in BMI could be accounted for by direct genetic effects based on a detailed analysis of genome-wide SNP data. We analyzed paternal BMI obtained from maternal ratings of their partner’s weight and height in 7829 trios. If any maternal biases are inherent in these ratings, including an indirect maternal genetic effect may allow us to still obtain valid estimates of the contributions from direct genetic effects.

A box-cox transformation and a scaling to zero mean and unit variance was applied to all phenotype measures. We included gender as a covariate in analyses of offspring phenotypes. All models were fit using the OpenMx package (Version 2.13; Neale et al., 2016) in R (Version 3.6.0; R Core Team, 2014). R code for fitting the model can be found in the supplementary material.

Results from applying the full model to the four phenotypes are presented in table 1. For all phenotypes except maternal relationship satisfaction, direct effects accounted for the largest portion of genetic influences. For birth weight and temperament, direct effects are estimated as the offspring genetic effects 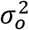, accounting for 12% of the variance for both phenotypes. For maternal relationship satisfaction direct genetic effects 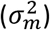 accounted for 8% of the variance, whereas for paternal BMI, direct genetic effects 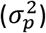 accounted for 32% of the variance.

**Table 1.** Parameter estimates and standard errors from the fitted models.

| Phenotype | Focal | $\sigma_m^2$ | $\sigma_p^2$ | $\sigma_o^2$ | $\sigma_{om}$ | $\sigma_{op}$ | $\sigma_{pm}$ | $\sigma_e^2$ |
| --- | --- | --- | --- | --- | --- | --- | --- | --- |
| Birth weight | Offspring | 0.08 (0.05) | 0.00* | 0.12 (0.06) | 0.01 (0.05) | -0.01 (0.03) | 0.05 (0.03) | 0.78 (0.05) |
| Temperament | Offspring | 0.04 (0.05) | 0.00* | 0.12 (0.07) | -0.01 (0.05) | -0.02 (0.04) | -0.01 (0.04) | 0.86 (0.05) |
| Relationship satisfaction | Mother | 0.08 (0.05) | 0.03 (0.06) | 0.10 (0.07) | -0.02 (0.05) | 0.01 (0.05) | 0.06 (0.04) | 0.81 (0.06) |
| BMI | Father | 0.04 (0.05) | 0.32 (0.05) | 0.00* | -0.02 (0.03) | -0.06 (0.04) | 0.07 (0.04) | 0.71 (0.05) |

The second column indicates the focal individual in the analysis; \* = Parameter is estimated at boundary and no standard error could be computed.

The strongest indications for indirect genetic effects were found for offspring birth weight - where 8% of the variability in birth weight could be ascribed to maternal genetic effects 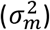 – and maternal relationship satisfaction, where 10% of the variability in relationship satisfaction could be ascribed to offspring genetic effects 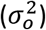 – a larger fraction than that due to direct genetic effects.

As the results indicate both direct and indirect genetic effects for offspring birth weight and maternal relationship satisfaction, it is of further interest to analyze the covariance between these components. The genetic correlation between maternal and offspring effects for birth weight was small, estimated as *σ*_*om*_/(*σ*_*m*_*σ*_*o*_) = 0.07. A stronger, negative genetic correlation of *σ*_*om*_/(*σ*_*m*_*σ*_*o*_) = −0.27, was estimated between maternal and offspring effects with respect to maternal reported relationship satisfaction.

Overall, these results are consistent with the general findings from twin studies, pointing to direct additive genetic effects as the major systematic source of variation for most traits (McAdams et al., 2014; Polderman et al., 2015). We also found indications for indirect genetic effects, most markedly for offspring birth weight and maternal relationship satisfaction. With respect to birth weight, these results are in line with previous findings. The weak genetic correlation further suggests that different genes may be involved in these effects. With respect to maternal relationship satisfaction, these findings are novel and may motivate further studies into the role of how relationship satisfaction may depend on infant characteristics. The negative genetic correlation may imply that some genes may have the opposite effects in mothers and offspring.

Considering the relatively large uncertainty associated with the parameter estimates, the results from the applications should be interpreted with caution. We emphasize that our analyses are not intended as a comprehensive study of the causes of variation for the phenotypes we examined, but rather are meant to illustrate how the proposed model can be used to investigate a diverse range of research questions.. Some of the solutions also gave meaningless results for the joint distributions of the genetic effects, with parameter estimates at the boundary or implied correlations greater than one. This is likely an indication that there is not enough data to support the model complexity, or that the model is misspecified. For a more detailed analysis it would likely be preferable to fit alternative nested models as described above, and test whether simpler models are equally supported by the data. Considerably larger sample sizes may be necessary to justify reliable inferences about the model parameters (Visscher et al., 2014; Yang, Zeng, Goddard, Wray, & Visscher, 2017). However, such sample sizes are increasingly available.

## Discussion

We proposed a new method, Trio-GCTA, for resolving direct and indirect genetic effects within parent-offspring trios when genome-wide SNP data is available. The model formulation is invariant to which of the family members is the focal individual in the analysis; only the interpretation of parameters (in terms of direct and indirect genetic effects) changes in different cases. We illustrated this by applying the method to four exemplar phenotypes using real data on offspring, maternal and paternal phenotypes. Results from the applications highlighted the potential of the method for clarifying intra-familial dynamics.

An advantage of the proposed method is the ability to gain insights into the dynamics of intra-familial processes without requiring specification of the specific traits that mediate the indirect genetic effects. Variance-partitioning of direct and indirect genetic effects may therefore serve as a useful first step, potentially motivating more detailed studies of specific processes. Trait-based models (Bijma, 2014), including explicit formulations of the hypothesized mediating variables may potentially provide better understanding of such mechanisms. However, in addition to the computational challenges, such specifications would also contradict one of the initial motivations for the GCTA model which avoid bias from common environmental effects by relying on measures obtained from unrelated individuals (Yang et al., 2011).

There are several issues related to estimating genetic variance parameters from genome-wide SNP data. Yang et al. (2017) emphasized that genetic variance parameters based on measured (or imputed) genome-wide SNPs differ from population parameters because they are dependent on the specific set of SNPs included in the analysis. They addressed several issues relating to estimating genetic variance parameters from genome-wide SNP data, and these considerations apply also to the method proposed in the current paper. There are likely further challenges that are specifically related to the use of parent-offspring trios and the method we have proposed here. First, the full model has seven variance parameters, which will likely require large sample sizes in order to obtain reliable estimates. Second, we have assumed that mating is random, and it is currently unclear how assortative mating could affect inferences under different models of intra-familial interactions. Third, although the distinction between direct and indirect genetic effects of parents and offspring may be an adequate description of many phenotypes, other relatives such as siblings may also play important roles in determining individual differences.

We believe the proposed method will provide a useful tool for researchers interested in the complexity of intra-familial dynamics, allowing investigations of research questions that may otherwise be difficult to study.

## Acknowledgements

The Norwegian Mother, Father and Child Cohort Study is supported by the Norwegian Ministry of Health and Care Services and the Ministry of Education and Research. We are grateful to all the participating families in Norway who take part in this on-going cohort study.

This publication is a part of the project “Intergenerational Transmission of Internalizing and Externalizing Psychopathological Spectra: A Genome-Wide Complex Trait Study” supported by the Research Council of Norway (262177). Eivind Ystrom was supported by the Norwegian Research Council (262177 and 288083). Laurie Hannigan was supported by a grant from the South-Eastern Norway Regional Health Authority (2018059). Alexandra Havdahl was supported by the South-Eastern Norway Regional Health Authority (2018058 and 2020022). Tom A. McAdams was supported by a Sir Henry Dale Fellowship, jointly funded by the Wellcome Trust and the Royal Society (107706/Z/15/Z) and the Norwegian Research Council (288083). Eshim S Jami was supported by the European Union’s Horizon 2020 research and innovation programme, Marie Sklodowska Curie Actions (721567).

We thank the Norwegian Institute of Public Health (NIPH) for generating high-quality genomic data. This research is part of the HARVEST collaboration, supported by the Research Council of Norway (#229624). We further thank the Center for Diabetes Research, the University of Bergen for providing genotype data and performing quality control and imputation of the data funded by the ERC AdG project SELECTionPREDISPOSED, Stiftelsen Kristian Gerhard Jebsen, Trond Mohn Foundation, the Research Council of Norway, the Novo Nordisk Foundation, the University of Bergen, and the Western Norway health Authorities (Helse Vest).

