## Supplementary file 1 for "Direct and indirect effects of maternal, paternal, and offspring genotypes: Trio-GCTA"

### Supplementary 1: R code for simulating data and fitting the model.

This document provides R code for simulating a dataset and fitting a model with direct and indirect genetic effects to parent-offspring trio data using OpenMx (Neale et al. 2016).

#### Genotypes

$K = 1000$ parent-offspring trios and $M = 300$ SNPs are simulated with genotypes depending on the outcome of two binomial trials with allele frequencies sampled uniformly from 0.4 to 0.6.

```
set.seed(080318)
library(OpenMx)
K = 1000
M = 300
P = runif(M, 0.4, 0.6) # Allele frequencies
# Maternal and paternal transmitted and untransmitted alleles
X_mt = matrix(rbinom(K * M, 1, P), K, M, byrow = T)
X_pt = matrix(rbinom(K * M, 1, P), K, M, byrow = T)
X_mu = matrix(rbinom(K * M, 1, P), K, M, byrow = T)
X_pu = matrix(rbinom(K * M, 1, P), K, M, byrow = T)
# Genotypes
X_m = X_mt + X_mu
X_p = X_pt + X_pu
X_o = X_mt + X_pt
# Standardized genotypes
Z_m = scale(X_m, 2 * P, sqrt(2 * P * (1 - P)))
Z_p = scale(X_p, 2 * P, sqrt(2 * P * (1 - P)))
Z_o = scale(X_o, 2 * P, sqrt(2 * P * (1 - P)))
```

#### Genetic, environmental and phenotypic values

We then sample correlated allelic effect sizes and independent environmental deviations from normal distributions. Phenotypes are functions of the standardized genotypes, allelic effects and environmental deviations.

```
# Allelic effects
Sigma_u = matrix(c(3, 0.7, 1, 0.7, 2, 0.5, 1, 0.5, 1.5), 3, 3)
B = MASS::mvrnorm(M, c(0, 0, 0), Sigma_u / M)
# Environmental deviations
e = rnorm(K)
# Phenotypes
y = cbind(Z_m, Z_p, Z_o) %*% c(B) + e
```

#### GRMs

The genetic relatedness matrices among individuals are computed as in Yang et al. (2011).

```r
A_mm = tcrossprod(Z_m) / M
A_pp = tcrossprod(Z_p) / M
A_oo = tcrossprod(Z_o) / M
A_pm = tcrossprod(Z_p, Z_m) / M
A_mp = tcrossprod(Z_m, Z_p) / M
A_om = tcrossprod(Z_o, Z_m) / M
A_mo = tcrossprod(Z_m, Z_o) / M
A_op = tcrossprod(Z_o, Z_p) / M
A_po = tcrossprod(Z_p, Z_o) / M
A_ommo = A_om + A_mo
A_oppo = A_op + A_po
A_pmmp = A_pm + A_mp
```

##### Fit the model with OpenMx

```r
y = scale(y, T, F)
colnames(y) = "y"
mod_mpo = mxModel("mod_mpo",
              # Data
              mxData(y, "raw", sort = F),
              mxMatrix("Sy", K, K, F, A_mm, name = "A_mm"),
              mxMatrix("Sy", K, K, F, A_pp, name = "A_pp"),
              mxMatrix("Sy", K, K, F, A_oo, name = "A_oo"),
              mxMatrix("Sy", K, K, F, A_ommo, name = "A_ommo"),
              mxMatrix("Sy", K, K, F, A_oppo, name = "A_oppo"),
              mxMatrix("Sy", K, K, F, A_pmmp, name = "A_pmmp"),
              mxMatrix("Id", K, K, name = "I"),
              # Parameters
              mxMatrix("Fu", 1, 1, T, Sigma_u[1, 1], "m2", name = "M_m2"),
              mxMatrix("Fu", 1, 1, T, Sigma_u[2, 2], "p2", name = "M_p2"),
              mxMatrix("Fu", 1, 1, T, Sigma_u[3, 3], "o2", name = "M_o2"),
              mxMatrix("Fu", 1, 1, T, Sigma_u[3, 1], "om", name = "M_om"),
              mxMatrix("Fu", 1, 1, T, Sigma_u[3, 2], "op", name = "M_op"),
              mxMatrix("Fu", 1, 1, T, Sigma_u[2, 1], "pm", name = "M_pm"),
              mxMatrix("Fu", 1, 1, T, 1, "e2", name = "M_e2"),
              # Covariance
              mxAlgebra(m2 * A_mm + p2 * A_pp + o2 * A_oo + e2 * I +
                          om * A_ommo + op * A_oppo + pm * A_pmmp, name = "V"),
              mxExpectationGREML("V", dataset.is.yX = T),
              mxFitFunctionGREML())
fit = mxRun(mod_mpo)
```

```
## Running mod_mpo with 7 parameters
```

```r
summary(fit)
```

```
## Summary of mod_mpo
##
## free parameters:
##   name matrix row col  Estimate  Std.Error A
## 1   m2   M_m2   1   1 3.0576426 0.33024097
## 2   p2   M_p2   1   1 1.8222523 0.23192279
## 3   o2   M_o2   1   1 1.6613206 0.24712446
```

```
## 4   om   M_om   1   1 0.9529161 0.20390213
## 5   op   M_op   1   1 0.5605705 0.16756934
## 6   pm   M_pm   1   1 0.5667008 0.20180056
## 7   e2   M_e2   1   1 0.9339887 0.08832124
##
## Model Statistics:
##                |  Parameters  |  Degrees of Freedom  |  Fit (-2lnL units)
##        Model:             7                    993              4250.19
##    Saturated:            NA                     NA                   NA
## Independence:            NA                     NA                   NA
## Number of observations/statistics: 1/1000
##
## Information Criteria:
##       |  df Penalty  |  Parameters Penalty  |  Sample-Size Adjusted
## AIC:       2264.19               4264.19                 4248.190
## BIC:       4250.19               4250.19                 4235.633
## CFI: NA
## TLI: 1   (also known as NNFI)
## RMSEA:  0  [95% CI (NA, NA)]
## Prob(RMSEA <= 0.05): NA
## To get additional fit indices, see help(mxRefModels)
## timestamp: 2020-03-19 21:36:00
## Wall clock time: 173.8285 secs
## optimizer:  CSOLNP
## OpenMx version number: 2.15.5
## Need help?  See help(mxSummary)
```

The model appear to have converged with reasonable parameter estimates.

#### Session information

```
## R version 3.6.2 (2019-12-12)
## Platform: x86_64-w64-mingw32/x64 (64-bit)
## Running under: Windows 10 x64 (build 17134)
##
## Matrix products: default
##
## locale:
## [1] LC_COLLATE=Norwegian Bokmål_Norway.1252
## [2] LC_CTYPE=Norwegian Bokmål_Norway.1252
## [3] LC_MONETARY=Norwegian Bokmål_Norway.1252
## [4] LC_NUMERIC=C
## [5] LC_TIME=Norwegian Bokmål_Norway.1252
##
## attached base packages:
## [1] stats     graphics  grDevices utils     datasets  methods   base
##
## other attached packages:
## [1] OpenMx_2.15.5
##
## loaded via a namespace (and not attached):
##  [1] Rcpp_1.0.3      lattice_0.20-38 digest_0.6.23   MASS_7.3-51.4
##  [5] grid_3.6.2      magrittr_1.5    evaluate_0.14   rlang_0.4.2
##  [9] stringi_1.4.3   Matrix_1.2-18   rmarkdown_2.0   tools_3.6.2
```

```
## [13] stringr_1.4.0   xfun_0.11       yaml_2.2.0      parallel_3.6.2
## [17] compiler_3.6.2  htmltools_0.4.0 knitr_1.26
```
