## Supplementary file 1 for "Direct and indirect effects of maternal, paternal, and offspring genotypes: Trio-GCTA"

### Supplementary 2: Simulation study

We performed a small simulation study in order verify that the model is able to recover population parameters under ideal conditions.

#### Design

We generated data from $K = 1000$ parent-offspring trios and $M = 300$ SNPs with direct and indirect effects. Maternal and paternal genotypes were simulated as the outcome of two binomial trials with allele frequencies sampled uniformly from 0.4 to 0.6. Offspring genotypes were generated by summing one allele from each parent. Allelic effects were sampled from a normal distribution

$$\begin{bmatrix} \boldsymbol{u}_m \\ \boldsymbol{u}_p \\ \boldsymbol{u}_o \end{bmatrix} \sim \mathcal{N}\left( \begin{bmatrix} \mathbf{0} \\ \mathbf{0} \\ \mathbf{0} \end{bmatrix}, \begin{bmatrix} \frac{3}{M}\mathbf{I} & \frac{0.7}{M}\mathbf{I} & \frac{1}{M}\mathbf{I} \\ \frac{0.7}{M}\mathbf{I} & \frac{2}{M}\mathbf{I} & \frac{0.5}{M}\mathbf{I} \\ \frac{1}{M}\mathbf{I} & \frac{0.5}{M}\mathbf{I} & \frac{1.5}{M}\mathbf{I} \end{bmatrix} \right).$$

Environmental deviations were sampled from $\boldsymbol{e} \sim \mathcal{N}(\mathbf{0}, \mathbf{I})$. The genotypes was then standardized and phenotypes were generated as a linear combination of the standardized genotypes, allelic effects and environmental deviations

$$\boldsymbol{y} = \mathbf{Z}_m\boldsymbol{u}_m + \mathbf{Z}_p\boldsymbol{u}_p + \mathbf{Z}_o\boldsymbol{u}_o + \boldsymbol{e}.$$

We repeated the same procedure for 1000 repetitions and fitted the model described in the main paper.

#### Results

Table 1 displays results from the simulation study. We examined bias and variability in parameter estimates using the formulas provided in Morris, White and Crowther (2019). For all parameters, the average of all simulation estimates were close to the simulation values. The estimated bias was less than 1% for all parameters.

Table 1: Summary from simulation study.

| | Expected | Mean | Bias | Bias (%) | SD | SE |
|---|---|---|---|---|---|---|
| $\sigma_m^2$ | 3.0 | 3.001 | 0.001 | 0.046 | 0.335 | 0.011 |
| $\sigma_p^2$ | 2.0 | 1.992 | -0.008 | -0.384 | 0.246 | 0.008 |
| $\sigma_o^2$ | 1.5 | 1.490 | -0.010 | -0.637 | 0.240 | 0.008 |
| $\sigma_{om}$ | 1.0 | 1.003 | 0.003 | 0.306 | 0.192 | 0.006 |
| $\sigma_{op}$ | 0.5 | 0.503 | 0.003 | 0.545 | 0.172 | 0.005 |
| $\sigma_{pm}$ | 0.7 | 0.693 | -0.007 | -0.962 | 0.219 | 0.007 |
| $\sigma_e^2$ | 1.0 | 1.007 | 0.007 | 0.746 | 0.094 | 0.003 |

The points in figure 1 displays mean bias estimated across the simulations. The vertical lines indicates two times the standard error above and below the estimates. The deviations from zero appears to be largely consistent with sampling error.

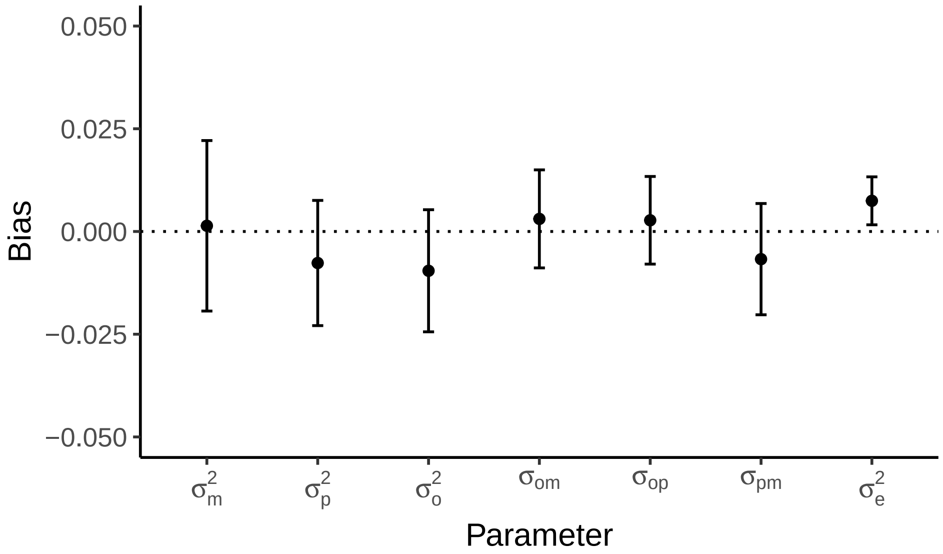


Figure 1: Average bias across simulations.
